# ONC201/TIC10 enhances durability of mTOR inhibitor everolimus in metastatic ER+ breast cancer

**DOI:** 10.1101/2022.12.27.522019

**Authors:** Elena Farmaki, Aritro Nath, Rena Emond, Kimya L Karimi, Vince K Grolmusz, Patrick A Cosgrove, Andrea H Bild

## Abstract

The mTOR inhibitor, everolimus, is an important clinical management component of metastatic ER+ breast cancer. However, most patients develop resistance and progress on therapy, highlighting the need to discover strategies that increase mTOR inhibitor effectiveness. We developed ER+ breast cancer cell lines, sensitive or resistant to everolimus, and discovered that combination treatment of ONC201/TIC10 with everolimus inhibited cell growth in 2D/3D *in vitro* studies. We confirmed increased therapeutic response in primary patient cells progressing on everolimus, supporting clinical relevance. We show ONC201/TIC10, in metastatic ER+ breast cancer cells, mechanistically involves oxidative phosphorylation inhibition and stress response activation. Transcriptomic analysis in everolimus resistant breast patient tumors and mitochondrial functional assays in resistant cell lines demonstrated increased mitochondrial respiration dependency, contributing to ONC201/TIC10 sensitivity. We propose that ONC201/TIC10 and modulation of mitochondrial function may provide an effective add-on therapy strategy for patients with metastatic ER+ breast cancers resistant to mTOR inhibitors.

## Introduction

Breast cancer (BC) is a fatal disease with 287,850 new cases in 2022 in the United States and an estimated mortality rate of 43,250 (https://seer.cancer.gov/statfacts/html/breast.html). The most common BC subtype is positive for hormone receptor (HR+) and does not overexpress the human epidermal growth factor receptor 2 (HER2/neu), accounting for 68% of all breast cancer cases (https://seer.cancer.gov/statfacts/html/breast-subtypes.html). The primary treatment for estrogen receptor positive (ER+), HER2-breast cancer are therapies targeting estrogen signaling; however, resistance to these therapies in metastatic BC (mBC) and disease progression remains a significant challenge (Osborne and Schiff, 2011).

Aberrant activation of the mammalian (mechanistic) target of rapamycin (mTOR) signaling has been identified in multiple human tumors (Hare and Harvey, 2017; Ortolani et al., 2015) and is involved in resistance to endocrine therapy (Johnston, 2015; Osborne and Schiff, 2011). Everolimus, an analog of rapamycin, is an allosteric inhibitor of mTOR complex 1 (mTOR1), and in combination with the aromatase inhibitor exemestane improved progression-free survival in patients with advanced breast cancer (Baselga et al., 2012). Based on these clinical studies the combination was approved in 2012 as a second-line therapy for recurrent or metastatic breast cancer (National Comprehensive Cancer Network, NCCN guidelines) (Baselga et al., 2012). Despite its efficacy, a large subset of patients develops progression and resistance to this combination, highlighting the need for more effective therapeutic strategies for advanced breast cancer.

Studies on everolimus resistance in breast cancer revealed survival mechanisms including activation of AKT (Carew et al., 2011; Chen et al., 2013) and MAPK signaling (Campone et al., 2021; Carew et al., 2011; Kimura et al., 2018; Mendoza et al., 2011), upregulation of MYC (Bihani et al., 2015; Lui et al., 2016), increased oxidative stress (Neklesa and Davis, 2008), metabolic reprogramming (Pusapati et al., 2016), increased expression of anti-apoptotic molecules such as survivin (Taglieri et al., 2017), and activation of epithelial and mesenchymal transition (EMT) (Holder et al., 2015).

ONC201/TIC10 belongs to the imipridone class of inhibitors and is currently being evaluated in clinical trials for solid tumors including breast and endometrial cancer, gliomas, and hematological malignancies (Prabhu et al., 2020). The mechanism of ONC201/TIC10 action involves binding and activation of the mitochondrial protease caseinolytic protease P (ClpP) (Graves et al., 2019) and inhibition of the G protein coupled receptor (GPCR) dopamine receptor D2 (DRD2) (Kline et al., 2018). ONC201/TIC10 has been shown to synergize with everolimus in prostate cancer (Lev et al., 2018). Furthermore, colorectal cancer cells were more sensitive to the combination of ONC201/TIC10 with mTOR inhibitor AZD-8055 (Jin et al., 2016). While, the effectiveness of the combination of everolimus with ONC201/TIC10 has not been studied in breast cancer, upregulation of mitochondrial ClpP that is a target of ONC201/TIC10 has been reported both in breast cancer cells and tissues (Cormio et al., 2021; Luo et al., 2020). In addition, ONC201/TIC10 inhibits mitochondrial oxidative phosphorylation in breast cancer cells, causing depletion of cellular ATP (Dwucet et al., 2021; Greer et al., 2018; Ishida et al., 2018; Pruss et al., 2020). Increased dependancy on oxidative phosphorylation (OXPHOS) has been shown in chemotherapy resistant triple negative breast cancer (Evans et al., 2021). Therefore, targeting ClpP and mitochondrial metabolism could constitute a valid approach for increasing the efficacy of everolimus.

The aim of our study was to explore treatment strategies that can enhance the anti-tumor activity of standard of care therapy by targeting common resistant states specific to ER+ mBC. For our study, we developed *in vitro* models of acquired everolimus resistance for paired ER+ BC cell lines that are either sensitive or resistant to everolimus, and viable primary patient ER+ BC cells with known response to everolimus therapy. We used these models to identify therapies effective in everolimus resistant cancer cells and found that the small molecule ONC201/TIC10 can inhibit the proliferation of resistant cells by disrupting mitochondrial function and metabolism. RNA-sequencing analysis in ER+ mBC tumors non-responsive to everolimus and mitochondrial functional assays in resistant cell lines demonstrated increased dependency on mitochondrial respiration, further supporting the sensitivity to ONC201/TIC10. Thus, we demonstrate that ONC201/TIC10 enhances durability of everolimus in resistant ER+ BC cell lines as well as primary cells from patients progressing on everolimus.

## Results

### ONC201/TIC10 inhibits the proliferation of breast cancer cell lines sensitive and resistant to everolimus

Everolimus resistant cell lines were generated by long-term culture of parental cell lines in the continuous presence of 100nM everolimus for MCF7 and T47D or 50nM everolimus for CAMA-1 for 6-12 months (McDermott et al., 2014). Parental untreated cell lines were maintained in culture for the same time as the everolimus-treated cell lines as paired everolimus sensitive cell lines. To quantify the extent of resistance, we performed dose-response assays with increasing concentrations of everolimus in sensitive and everolimus resistant cells. Resistant cells showed a statistically significant reduced response to everolimus compared to sensitive cells (p<0.0001 for all cell lines, Figure 1A).

**Figure 1.**
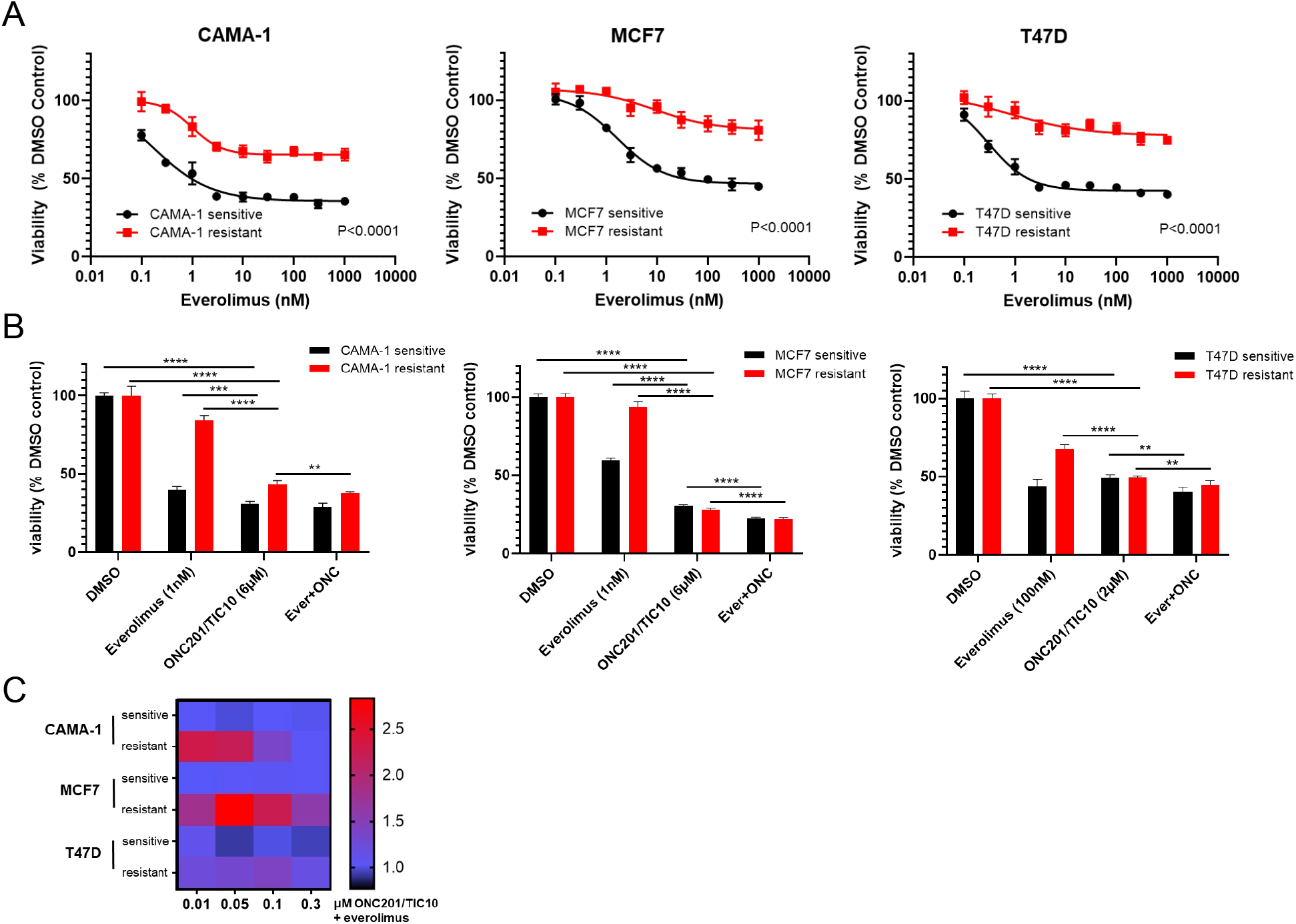
ONC201/TIC10 inhibits the proliferation of everolimus sensitive and resistant cells in 2D. A. Dose-response curves of CAMA-1, MCF7, and T47D sensitive and everolimus resistant cells under everolimus treatment. Cells were treated with increasing concentration of everolimus for 4 days and viability was measured using CellTiterGlo Chemiluminescent kit. B. Cell viability after 4 days treatment with DMSO, everolimus, ONC201/TIC10, or combination at the indicated concentrations. Data represents % viable cells compared with DMSO control treatment for each cell line and are shown as average of four replicates ± SD *, P < 0.05; **, P < 0.01; ***, P < 0.001, ****, P < 0.0001. C. Analysis of ONC201/TIC10 and everolimus interactions in 2D. Cells were treated with 1nM everolimus (CAMA-1 and T47D) or 100nM everolimus (MCF7), ONC201/TIC10, or combination at the indicated concentrations for 4 days in 2D and viability was measured using CellTiterGlo Chemiluminescent kit. The average Bliss Interaction Index was calculated and plotted as a heatmap in which red represents synergy or and blue represents additivity.

Next, we assessed the anti-proliferative efficacy of ONC201/TIC10 in the everolimus sensitive and resistant cell lines using both 2-dimensional (2D) and 3-dimensional (3D) assays. Sensitive and resistant cells were labeled with lentivirus to express a fluorescent protein for monitoring each population’s growth when cultured (Venus and mCherry fluorescence for the sensitive and resistant cells, respectively). For the 2D assays, everolimus sensitive and resistant cells were treated with everolimus, ONC201/TIC10, or combination for 4 days. For the 3D assays everolimus sensitive or resistant spheroids were cultured in the presence of drug treatments for 18 days with media and drug replacement every 3-4 days. Images and fluorescence signal intensity measurements were also captured on days of media and drug replacement to monitor growth inhibition over time.

In the 2D assays, ONC201/TIC10 single agent inhibited the proliferation of CAMA-1, MCF7 and T47D sensitive and resistant cells in a dose-dependent manner with IC50 values ranging from 1.43-1.90 μM (Supplementary Figure 1). ONC201/TIC10 single agent treatment resulted in significant growth inhibition compared to control and everolimus single agent in the all the resistant cell lines and in the CAMA-1 and MCF7 sensitive cells (Figure 1B). Combination of ONC201/TIC10 with everolimus significantly decreased cell proliferation compared to ONC201/TIC10 single agent, in all the resistant cells and in the MCF7 and T47D sensitive cells, by approximately 3% decrease for CAMA-1 sensitive, 5.5% for CAMA-1 resistant, 8% for MCF7 sensitive, 5% for MCF7 resistant, 8.8% for T74D sensitive, 4.7% for T47D resistant (Figure 1B). Analysis of the drug interactions using the Bliss Interaction Index showed additive effect in the sensitive and T47D resistant cells and synergistic effect in the CAMA-1 and MCF7 resistant cells (Figure 1C).

In the 3D assays, ONC201/TIC10 single agent treatment resulted in significant growth inhibition compared to control in the all the resistant cell lines and in the T47D sensitive cells as indicated by the fluorescence intensity (Figure 2A and 2B). Even though everolimus single agent had increased inhibitory effect compared to ONC201/TIC10 single agent, the combination significantly inhibited spheroid growth in sensitive and resistant cell lines compared to both everolimus and ONC201/TIC10 single agent (Figure 2A and 2B). Drug interactions analysis showed moderately better than additive effect in the CAMA-1 resistant cells as well as in T47D sensitive and resistant cells (Figure 2C).

**Figure 2.**
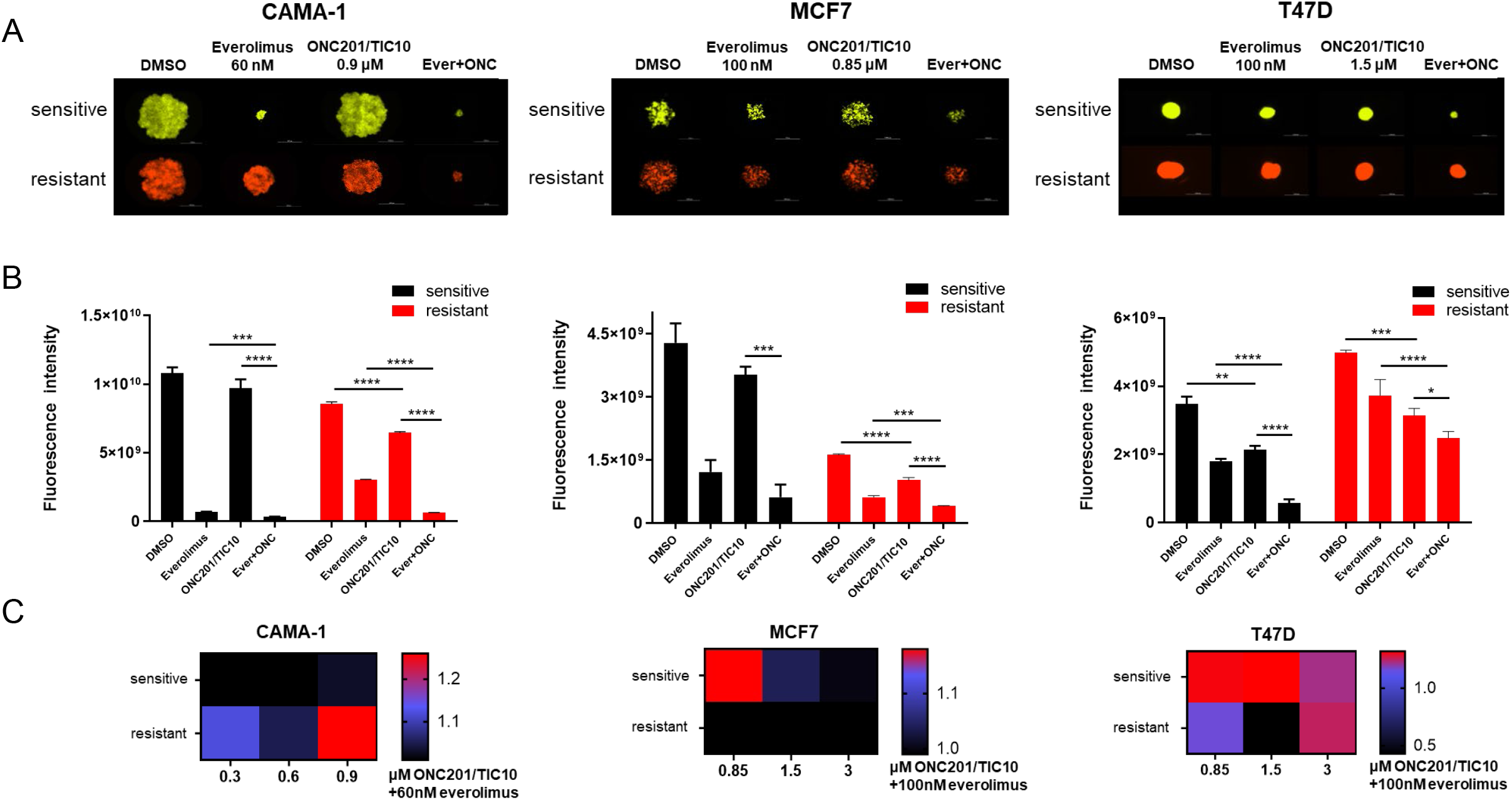
Combination of ONC201/TIC10 and everolimus inhibits spheroid growth in 3D. A. Representative images of spheroid growth of sensitive (Venus, green) or resistant (mCherry, red) cells cultured in everolimus, ONC201/TIC10 or combination treated media at the indicated concentrations for up to 18 days. B. Fluorescence intensity of sensitive and resistant cells under various treatment conditions. Data are represented as average of three replicates ± SD *, P < 0.05; **, P < 0.01; ***, P < 0.001, ****, P < 0.0001. C. Analysis of ONC201/TIC10 and everolimus interactions in 3D. Spheroids were cultured in the presence of drug treatments for 18 days and fluorescence intensity measurements were captured. The average Bliss Interaction Index was calculated and plotted as a heatmap in which red represents synergy or and blue represents additivity.

### Combination therapy of ONC201/TIC10 and everolimus inhibits the growth of primary patient-derived ER+ BC cell spheroids

To further validate the effect of the combination of ONC201/TIC10 with everolimus in cells from patients’ tumors, we performed 3D assays with primary cells from pleural effusions or ascites from patients’ tumor samples. We used short-term experiments, compared to sensitive and resistant cell lines, since primary cells were more sensitive to the drug treatments. Patient samples were selected based on the treatment history, specifically refractory tumors from patients who had received hormonal therapy followed by several lines of treatment including everolimus and progressed (Figure 3A). Spheroids of patient-derived BC cells illustrate more accurately the characteristics of tumors *in vivo* and provide a more suitable model for the assessment of drug treatments compared with the 2D cultures (Imamura et al., 2015; Langhans, 2018). Spheroids from 5 different patients were treated with ONC201/TIC10, everolimus or combination for 4 days. The effect of drug treatment on growth inhibition was assessed by measurement of cell proliferation (Figure 3B). Our results indicate that combination therapy of ONC201/TIC10 with everolimus inhibits spheroid growth compared to everolimus single agent or ONC201/TIC10 single agent (ONC201/TIC10 single agent versus combination 12.5% cell proliferation decrease for P1, 11% for P2, 4% for P3, 0.2% for P4 and 10.5% for P5) (Figure 3B). Importantly, we compared the response of patient’s cells during everolimus treatment and after everolimus resistance, based on the availability of samples for P2 that were collected over the course of treatment. We used these two different tumor samples, from the same patient, during everolimus treatment or post-everolimus and during chemotherapy treatment (Figure 3C). ONC201/TIC10 single agent or combination of ONC201/TIC10 with everolimus exhibited greater growth inhibition in the post-everolimus treatment cells (Figure 3C, Supplementary Figure 2) as shown by the significant reduction in proliferation of these cells compared to the cells acquired during everolimus treatment (7.2% cell decrease in cell proliferation during everolimus vs post-everolimus with ONC201/TIC10 and 9.2% cell decrease in cell proliferation during everolimus vs post-everolimus with the combination) (Figure 3C) and decreased spheroid size (Supplementary Figure 2). These results highlight the efficacy of ONC201/TIC10 on the patient-derived resistant cells.

**Figure 3.**
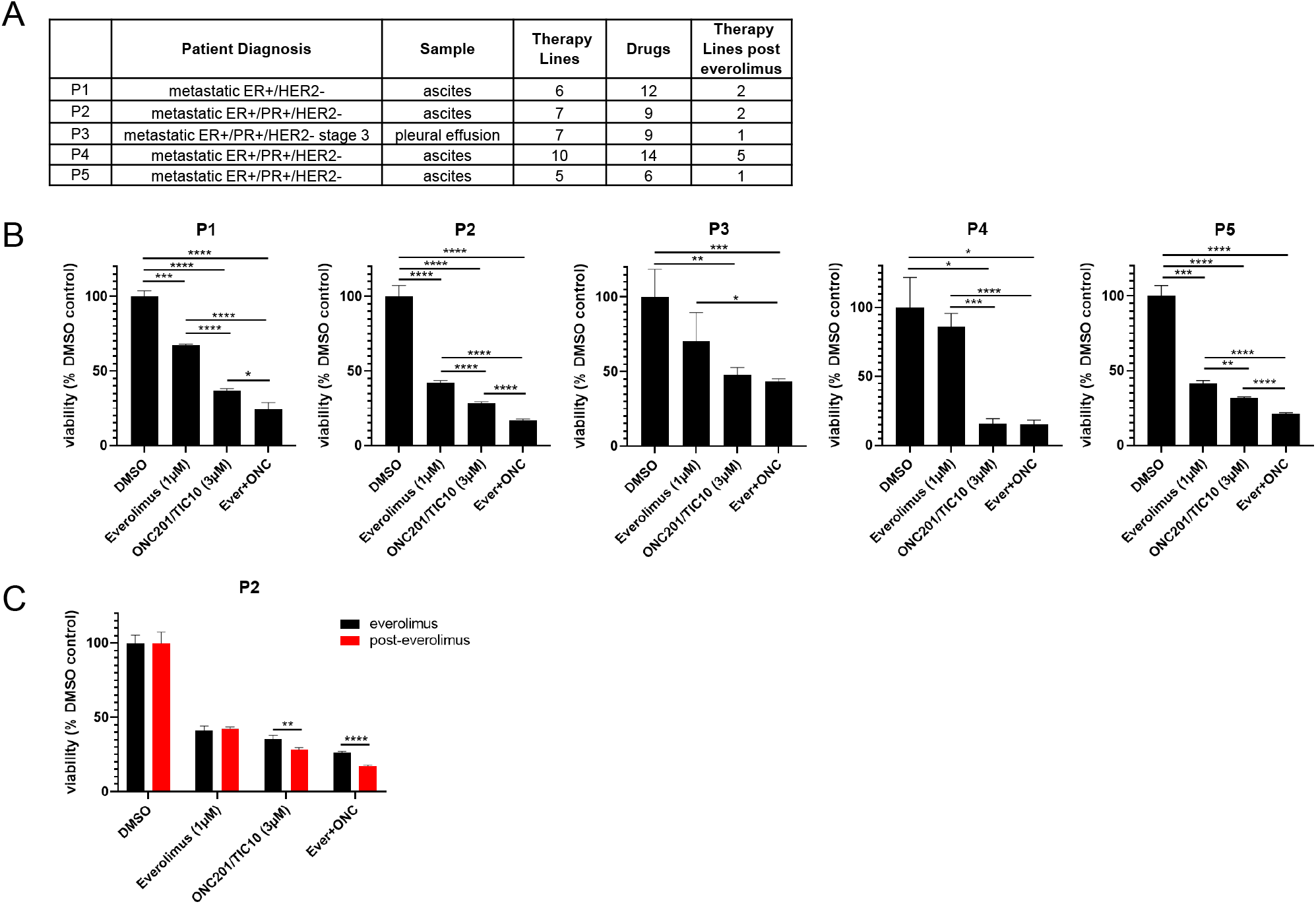
Combination therapy of ONC201/TIC10 and everolimus inhibits the growth of primary patient-derived cell spheroids. A. History of patients included in the study. B. 3D cultures of primary patient-derived ER+ BC cells from ascites or pleural effusions treated with everolimus, ONC201/TIC10 or combination at the indicated concentrations for 4 days. Cell viability was measured using CellTiterGlo Chemiluminescent kit. C. 3D cultures of primary patient-derived ER+ BC cells, while on everolimus treatment or post-everolimus treatment. Spheroids were treated with everolimus, ONC201/TIC10 or combination at the indicated concentrations for 4 days. Data represents % viable cells compared with DMSO control treatment and are shown as average of three replicates ± SD *, P < 0.05; **, P < 0.01; ***, P < 0.001, ****, P < 0.0001.

### ONC201/TIC10 causes loss of mitochondrial proteins and activation of stress response in everolimus sensitive and resistant cells

To identify the mechanism of ONC201/TIC10 sensitivity in the sensitive and resistant cells, we investigated the effects of the treatments on downstream signaling pathways, including mitochondrial pathways and stress pathways that have been previously described for BC cells under ONC201/TIC10 treatment (Greer et al., 2018; Ralff et al., 2017; Yuan et al., 2017). Western blot analysis showed that ONC201/TIC10 suppresses the protein expression of the mitochondrial proteins TFAM (mitochondrial transcription factor A) and TUFM (translation elongation factor Tu) in a dose-dependent manner in CAMA-1 and T47D sensitive and resistant cells (Figure 4A). TFAM and TUFM are important regulators of mitochondrial functions and are associated with the ONC201/TIC10-induced mitochondrial disruption (Graves et al., 2019; Greer et al., 2018).

**Figure 4.**
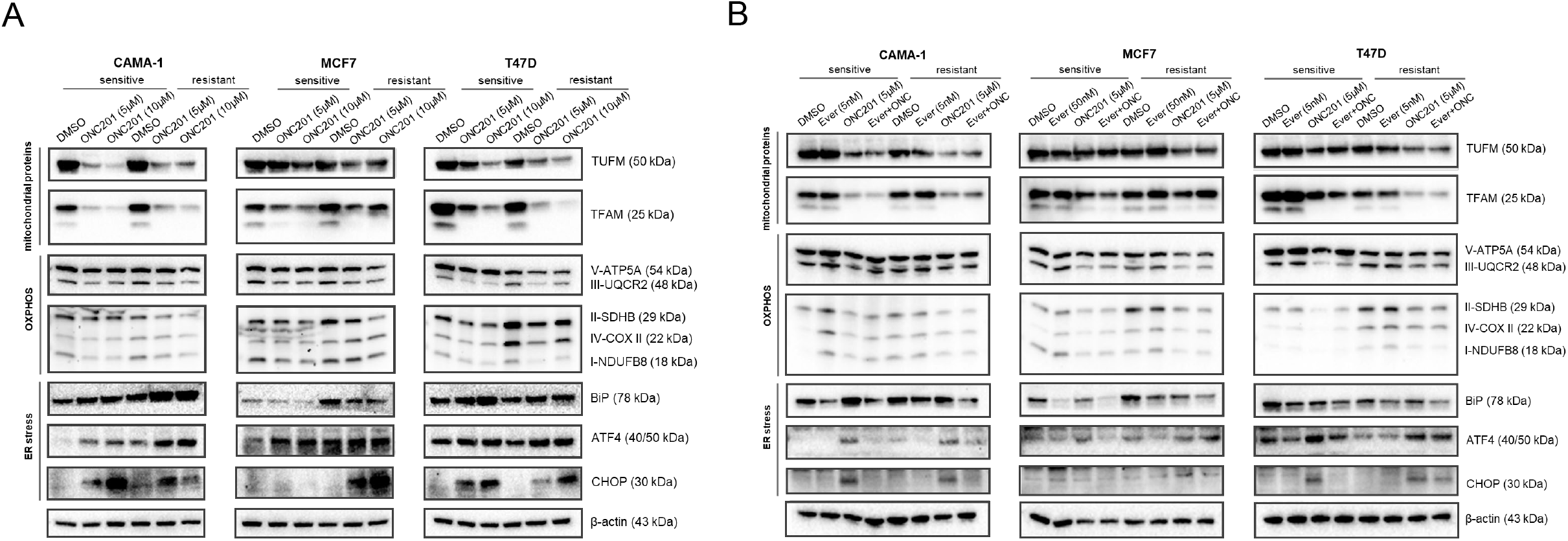
ONC201/TIC10 causes loss of mitochondrial proteins and activation of stress response in everolimus sensitive and resistant cells. A. CAMA-1, MCF7, T47D sensitive and everolimus resistant cells were treated for 24 hours with ONC201/TIC10 at the indicated concentrations, and cell lysates were immunoblotted for TUFM, TFAM, OXPHOS complexes (Complex I subunit NDUFB8, Complex II subunit 30kDa, Complex III subunit Core 2, Complex IV subunit II, and ATP synthase subunit alpha), BiP, ATF4, CHOP and β-actin. B. CAMA-1, MCF7, T47D sensitive and everolimus resistant cells were treated for 24 hours with everolimus, ONC201/TIC10 and combination at the indicated concentrations, and cell lysates were immunoblotted for the same proteins as above.

We further assessed the effect of ONC201/TIC10 on the oxidative phosphorylation proteins since ONC201/TIC10 toxicity has been linked to impaired OXPHOS in breast and brain tumors (Dwucet et al., 2021; Greer et al., 2018; Ishida et al., 2018; Pruss et al., 2020). ONC201/TIC10 suppresses the expression of the respiratory chain complexes in sensitive and resistant cells, specifically Complex I subunit NDUFB8, Complex II subunit 30kDa (CAMA-1 and MCF7) Complex III subunit Core 2 and ATP synthase subunit alpha (Figure 4A).

Next, we evaluated in our model the activation of the Integrated Stress Response (ISR), that has been reported to follow mitochondrial dysfunction in breast (Greer et al., 2018; Ralff et al., 2017; Yuan et al., 2017) and other malignancies (Al Madhoun et al., 2021; Fan et al., 2022; Lev et al., 2018). ONC201/TIC10 induces the expression of the chaperone BiP (HP70) and the pro-apoptotic transcription factors ATF4 and CHOP in a dose dependent-manner in CAMA-1 and T47D sensitive and the resistant cells (Figure 4A). Increased protein levels of CHOP were observed only in the MCF7 resistant cells as well as no increase in BiP levels suggesting the involvement of other UPR markers in these cells (Figure 4A).

Combination treatment of ONC201/TIC10 with everolimus had the same effect on the suppression of the mitochondrial proteins TFAM and TUFM and the OXPHOS complexes as the single agent (Figure 4B). Everolimus single agent did not affect the mitochondrial protein levels, as expected (Figure 4B). Activation of stress response was not observed with the combination in the sensitive cells, suggesting that everolimus alleviates endoplasmic reticulum stress through the activation of different cell death mechanisms (Figure 4B).

Inhibition of AKT, ERK and FoxO3a phosphorylation and activation of TRAIL that were previously described for ONC201/TIC10 (Allen et al., 2013; Ralff et al., 2017) were not observed in our model system (Supplementary Figure 3). Furthermore, ONC201/TIC10 inhibits the phosphorylation of ribosomal protein S6 that is downstream of mTOR signaling and this inhibition is increased in the resistant cell lines (Supplementary Figure 3).

To corroborate the mechanism of ONC201/TIC10 action observed in our experiments, we analyzed time course gene expression data of the MDA-MB-231 BC cell line treated with ONC201/TIC10 (Greer et al., 2018). Using a generalized linear model, we investigated the changes in pathway activity levels of the cell line measured at 0, 3, 6, 12 and 24 hours after treatment. Over the time course, ONC201/TIC10 treatment significantly reduced cell cycle pathway activity in the cell line (P = 1.8 x 10^-6^) (Figure 5A). Concurrently, we found a significant increase in the ATF4 target genes activated in response to ER stress (P = 1.2 x 10^-4^) (Figure 5B) and the UPR pathway activity (P = 1.8 x 10^-4^) (Figure 5C). In contrast, neither of the ERK (P = 0.55) (Figure 5D) or AKT (P = 0.61) (Figure 5E) signaling pathways showed any change in activity over time. These results confirm that the observed activity of ONC201/TIC10 in BC cells is via induction of mitochondrial stress rather than ERK/AKT inactivation.

**Figure 5.**
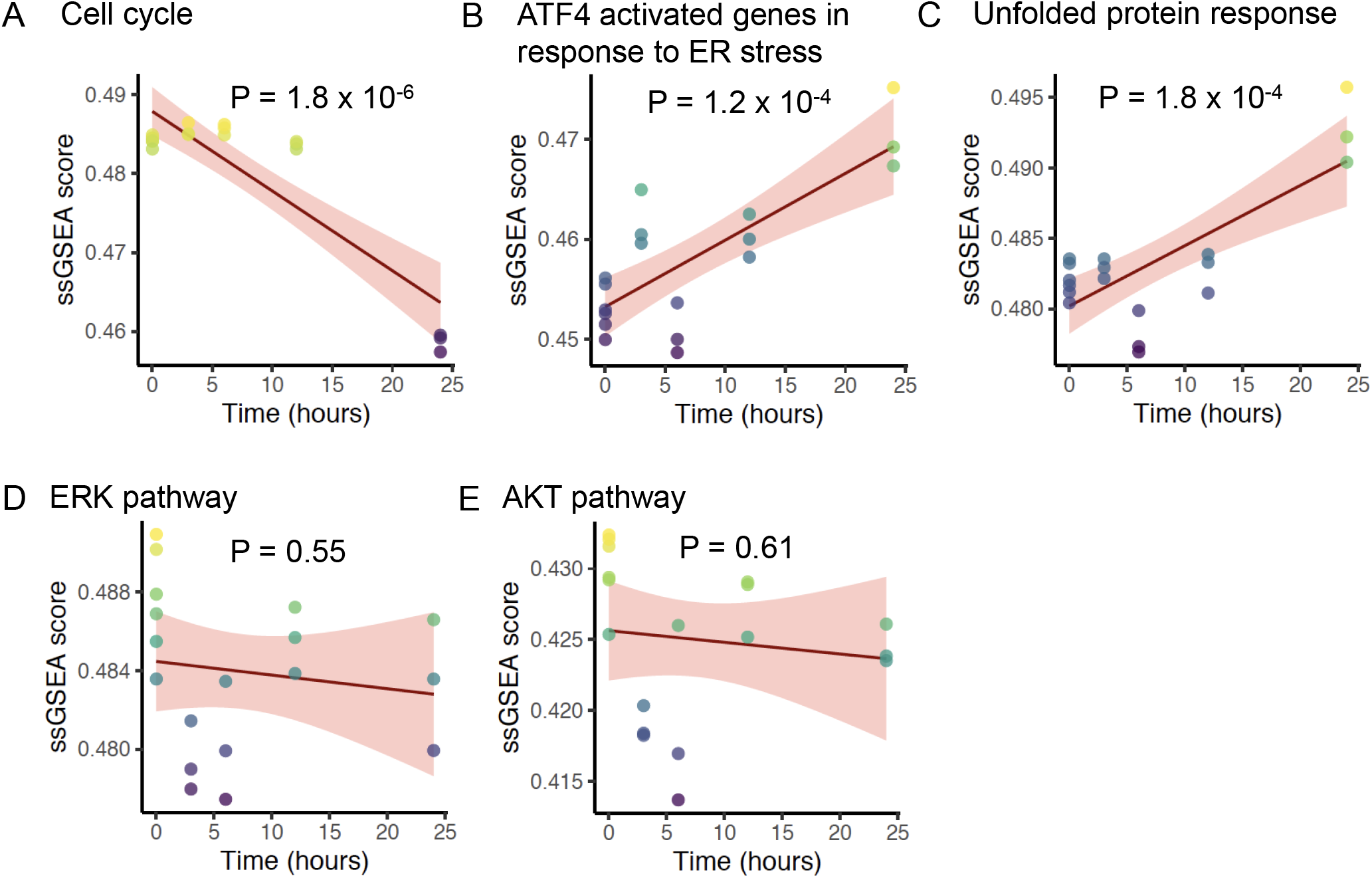
Change in pathway activity over time in response to ONC201/TIC10. Scatter plots displaying the enrichment scores (Y-axis) of A. REACTOME cell cycle signature, B. REACTOME ATF4 activates genes in response to endoplasmic reticulum stress signature, C. REACTOME unfolded protein response UPR signature, D. BIOCARTA ERK pathway signature, and E. BIOCARTA AKT pathway signature over time (X-axis). The solid lines and shaded area indicate linear fit and 95% confidence intervals respectively, with P-value of the fit indicated above each plot. The analysis includes data from six replicates at time 0 hour, and three replicates each at time 3, 6, 12 and 24 hours for a total n = 18.

### ONC201/TIC10 inhibits mitochondrial respiration in everolimus sensitive and resistant cells

Based on the findings for the mitochondrial pathways, we then assessed the mitochondrial respiratory capacity of sensitive and resistant cell lines using extracellular flux analysis. Comparison of the mitochondrial respiration between the sensitive and resistant cells indicates that CAMA-1 resistant cells have lower basal and maximal respiratory capacity compared to the sensitive cells while MCF7 and T47D resistant cells have significantly increased baseline, maximal and ATP-linked respiration compared to the sensitive cells suggesting increased dependency on OXPHOS for ATP production (Supplementary Figure 4). Furthermore, all resistant lines showed significantly increased baseline and stressed extracellular acidification rate (ECAR) (Supplementary Figure 4). Treatment with ONC201/TIC10 resulted in a significant decrease in oxidative consumption rates (OCR) compared to control and everolimus single agent, in both sensitive and resistant cell lines at baseline, maximal and ATP-linked respiration (Figure 6, Supplementary Figure 5). In addition, combination treatment of ONC201/TIC10 with everolimus further reduced mitochondrial respiratory capacities in all sensitive and resistant cells, except CAMA-1 cells that ONC201/TIC10 alone has similar effect on OCR as the combination (Figure 6, Supplementary Figure 5).

**Figure 6.**
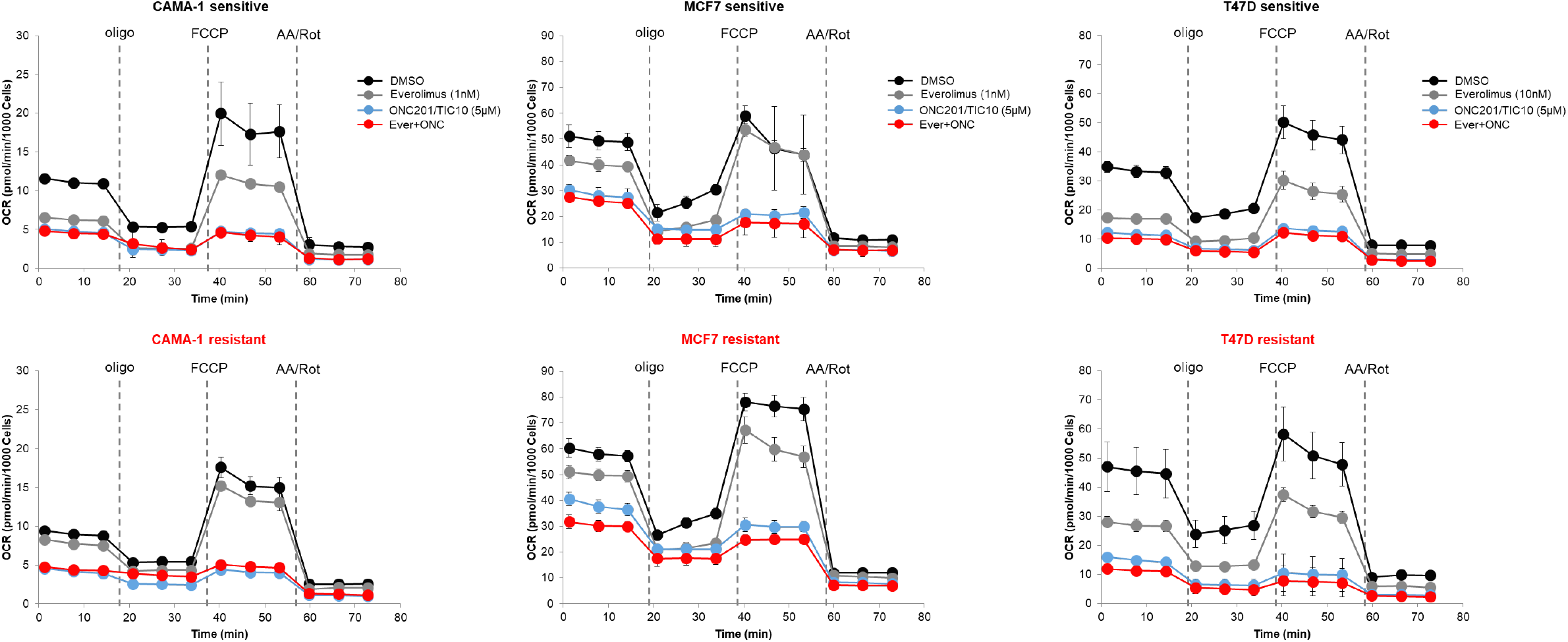
ONC201/TIC10 inhibits mitochondrial respiration in everolimus sensitive and resistant cells. Cells were treated for 18 hours with indicated concentrations of ONC201/TIC10 and everolimus. Mitochondrial respiration was measured using Seahorse XF Cell Mito Stress assay and oxygen consumption rates (OCR) are shown. Values were normalized to cell number generated from fluorescence intensity measurements and are represented as average of three replicates.

### Tumors non-responsive to everolimus sustain mitochondrial oxidative phosphorylation activity

We compared the changes in pathway phenotypes of mTOR inhibitor sensitive and resistant tumors using gene expression data from a neoadjuvant clinical trial of ER+ BC patients receiving everolimus (Sabine et al., 2010). We examined the changes in 21 patients by comparing single sample gene set enrichment scores of pathway signatures from the RNA-sequencing data from this trial taken before and after 11-14 days of treatment with everolimus. Patients classified as everolimus responders based on decrease in Ki67 staining post-treatment displayed significant decreased in cell cycle signature (P = 4.3 x 10^-3^), while non-responders did not show a significant change (P = 0.2) (Figure 7A). This result confirmed that everolimus failed to control the growth and proliferation of resistant tumors in an independent patient dataset. In the responder group, we found that everolimus treatment led to a significant decrease in the mitochondrial oxidative phosphorylation (P = 7.6 x 10^-3^), while there was no significant difference in the non-responders’ group (P = 0.83) (Figure 7B). These results suggest that non-responsive tumors continued to proliferate after everolimus treatment by utilizing oxidative phosphorylation pathway to fuel the growth of tumor cells, which is consistent with the findings from the resistant cell lines.

**Figure 7.**
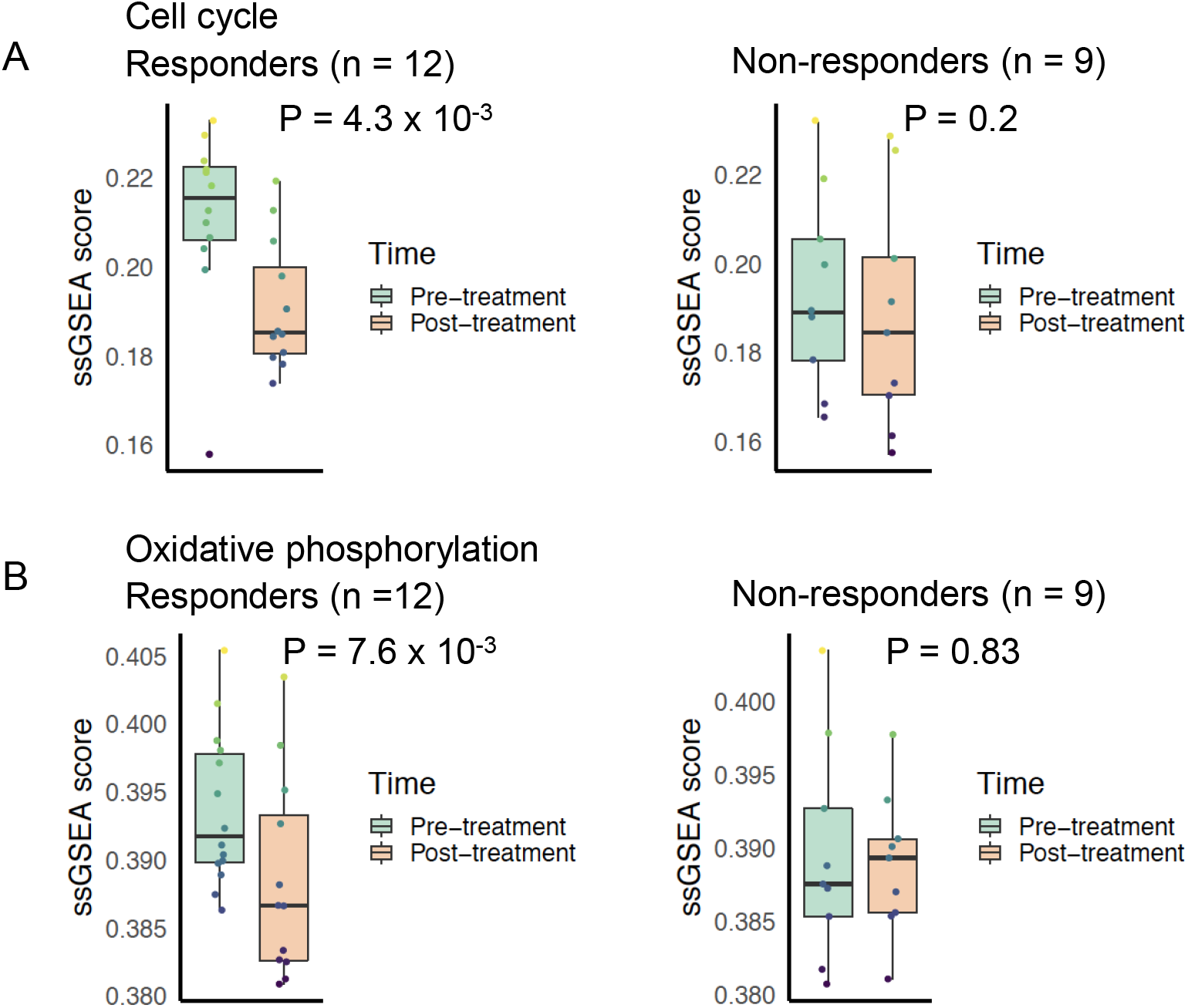
Cell cycle and oxidative phosphorylation pathway activity and neo-adjuvant everolimus response. Box plots comparing cell cycle pathway activity indicated by A. REACTOME cell cycle signature enrichment scores and B. KEGG oxidative phosphorylation signature enrichment scores between pre-treatment and post-treatment samples. The left panel shows patients that were classified as responders to everolimus, while the right panel shows patients that were classified as non-responders. Colored boxes indicate interquartile range, horizontal bars indicate median, and the whiskers indicate first and third quartiles. P-values from paired two-tailed t-test comparing pre-treatment vs. post-treatment scores are indicated above the plots.

## Discussion

Therapies targeting signaling pathways such as the CDK4/6 and the PI3K/AKT/mTOR pathway that have been identified among the pathways involved in endocrine therapy resistance (Johnston, 2015) have provided a significant clinical benefit in ER+ BC. The BOLERO-2 study showed that addition of everolimus to exemestane significantly improves progression-free survival, in post-menopausal ER+ BCs from 2.8 months on exemestane alone to 6.9 months on everolimus plus exemestane (Baselga et al., 2012). Despite this progress, resistance to targeted therapies still emerges in a substantial proportion of patients. Therefore, it is important to explore strategies to increase response to this therapy.

In the present study we explored treatment strategies to improve targeted therapy response and survival by combining drugs targeting resistance mechanisms. Specifically, we investigated if combination of everolimus with ONC201/TIC10 is effective for resistant BC cells and could provide a promising treatment strategy. Previous studies showed the efficacy of ONC201/TIC10 in TNBC cells (Greer et al., 2018; Yuan et al., 2017). We focused on the effect of ONC201/TIC10 on advanced ER+ BC cells including everolimus resistant cells and patient-derived spheroids resistant to endocrine therapy and everolimus.

Our results show that ONC201/TIC10 and modulation of mitochondrial function is effective in drug resistant cancer cells. Combination therapy with ONC201/TIC10 and everolimus showed increased growth inhibition in everolimus resistant BC cell lines as well as resistant patient tumor cells in 2D and 3D studies. In the 2D culture setting, ONC201/TIC10 was effective both as single agent and in combination with everolimus with similar potency in sensitive and resistant cells. In the 3D setting however, ONC201/TIC10 in combination with everolimus had a robust effect on growth inhibition in both resistant and sensitive cells, compared to ONC201/TIC10 single agent that had a minor effect. Differences in drug potency between the 2D and 3D cultures have been previously reported for BC cells and are associated with lower proliferation, increased apoptosis, microenvironmental conditions like hypoxia and reduced nutrients and altered signaling in the 3D setting (Burdett et al., 2010, Hamilton, 1998, Lovitt et al., 2015). These differences highlight the importance of utilizing more than one experimental model based on the hypotheses that are being addressed and given the limitations of each. Consistent with the resistant cell lines, the combination therapy had increased anti-proliferative activity in the resistant patient-derived spheroids. Collectively, these results recapitulate the increased sensitivity to the combination therapy and further support clinical relevance.

With regards to the mechanism of ONC201/TIC10 anti-proliferative activity, we found that ONC201/TIC10 causes mitochondrial dysfunction, which is consistent with previous findings for BC cells, including suppression of mitochondrial proteins, decreased OXPHOS and activation of stress response (Greer et al., 2018). We observed this effect not only in the sensitive cells but also in everolimus resistant cells, which supports the similarity in potency of ONC201/TIC10 single agent in everolimus sensitive and resistant cells in 2D assays. Importantly, data from RNA-sequencing in everolimus resistant patient tumors and from mitochondrial functional assays in resistant cell lines demonstrated increased dependency on mitochondrial respiration supporting the sensitivity to ONC201/TIC10. Therefore, ONC201/TIC10 could be a suitable candidate for the treatment everolimus-resistant tumors that depend on mitochondrial oxidative phosphorylation activity for sustained growth and proliferation.

Combination of ONC201/TIC10 with everolimus maintained the same level of mitochondrial and OXPHOS protein suppression as ONC201/TIC10 single agent. However, the activation of ISR after combination treatment was maintained only in the everolimus resistant cells and not the sensitive. These findings support the increased sensitivity of the combination in everolimus resistant cells and suggest as a mechanism of anti-proliferative activity the disruption of mitochondria and activation of stress response. Increased gene expression and protein levels of pro-apoptotic transcription factors CHOP and ATF4 has been reported for ONC201/TIC10 treatment (Greer et al., 2018; Ishizawa et al., 2016; Kline et al., 2016). Indeed, CHOP has been shown to induce apoptosis through the regulation of different anti-apoptotic and pro-apoptotic genes, including genes encoding the Bcl-2 family proteins, GADD34, endoplasmic reticulum oxidoreductin 1 (ERO1α), Tribbles-related protein 3 (TRB3), and DOC (Hu et al., 2018; Szegezdi et al., 2006). In addition, CHOP can downregulate the expressions of Bcl-2, Bcl-xL, and Mcl-1, and upregulate the expression of BIM, causing increased BAK and BAX expression. Bcl-2, Bcl-xL, and Mcl-1 have been shown to be downregulated by imipridones and ONC201/TIC10 (Al Madhoun et al., 2021; Rumman et al., 2021; Staley et al., 2021).

ATF4 can promote apoptosis either through regulating CHOP or independent of CHOP (Wortel et al., 2017). ATF4 also downregulates the anti-apoptotic Bcl-2 protein and upregulates pro-apoptotic signaling through the proteins BIM, NOXA, and PUMA (Szegezdi et al., 2006; Wortel et al., 2017). ONC201/TIC10 caused apoptosis through ATF4 in lymphoma and leukemia cells and inhibited mTORC1 signaling through the upregulation of ATF4 and DDIT4 (Ishizawa et al., 2016; Wang and Dougan, 2019).

Our results show that ISR was not activated upon combination treatment with ONC201/TIC10 and everolimus in sensitive cells, suggesting that sensitivity to everolimus in these cells can result in the activation of cell death mechanisms bypassing ISR. In line with this assumption, mTOR inhibitors are known to induce apoptosis through decreasing expression levels of various anti-apoptotic proteins, including Bcl-2, Bcl-xL, Mcl-1, and survivin (Du et al., 2018; Mills et al., 2008; Wangpaichitr et al., 2008). Other mechanisms of ONC201/TIC10 anti-proliferative activity in the everolimus resistant cells might include the involvement of c-Myc. High levels of c-Myc have been shown to be a predictive factor for growth inhibition and apoptosis by imipridones in glioblastoma (Ishida et al., 2018). In addition, the role of the Myc gene in promoting mTOR inhibitor resistance has been described in ER+ BC (Bihani et al., 2015). In conclusion, we used pre-clinical and clinical models to characterize sensitivity to mTOR inhibition. Combining experimental and clinical patient derived data with resistant cell line driven experiments, our study provides validated findings that are consistent across different contexts, thereby strengthening the potential for clinical applications of ONC201/TIC10. Collectively our findings suggest that ONC201/TIC10 could be used as an add-on treatment after mTOR therapy progression. Further *in vitro* as well as *in vivo* testing of this combination would be necessary to determine the optimal dosing and timing strategy for future clinical trials.

## Acknowledgments

This study was supported by a US National Cancer Institute award number U01CA264620 and U54CA209978 awarded to A.H.B.

## Author contributions

E.F: investigation, methodology, formal analysis, visualization, writing-original draft, writing–review and editing. A.N.: data curation, methodology, formal analysis, visualization, writing-original draft. R.E: investigation, methodology, formal analysis, visualization, writing-original draft. K.L.K: investigation, methodology, formal analysis. V.K.G: methodology, resources. P.A.C methodology, resources, editing draft. A.H.B. conceptualization, supervision, resources, funding acquisition, writing–review and editing.

---

**Supplementary Figure 1. ONC201/TIC10 inhibits the proliferation of everolimus sensitive and resistant cells.** Dose-response curves of CAMA-1, MCF7, and T47D sensitive and everolimus resistant cells under ONC201/TIC10 treatment. Cells were treated with increasing concentration of ONC201/TIC10 for 4 days and viability was measured using CellTiterGlo Chemiluminescent kit.

**Supplementary Figure 2. Combination therapy of ONC201/TIC10 and everolimus inhibits the growth of primary patient-derived cell spheroids.** Representative images of spheroid growth of primary patient-derived cells collected during everolimus treatment or post-everolimus and therapy resistant, treated with everolimus, ONC201/TIC10 or combination at the indicated concentrations for 4 days.

**Supplementary Figure 3. ONC201/TIC10 mechanism in everolimus sensitive and resistant is TRAIL-independent.** CAMA-1, MCF7, T47D sensitive and everolimus resistant cells were treated for 24 hours with everolimus, ONC201/TIC10 and combination at the indicated concentrations. Cell lysates were immunoblotted for pAKT, AKT, pERK, ERK, pFoxO3a, FOXO3, TRAIL, pS6, S6 and β-actin.

**Supplementary Figure 4. Mitochondrial respiration in everolimus sensitive and resistant cells.** Cells were cultured for 18 hours. Mitochondrial respiration was measured using Seahorse XF Cell Mito Stress assay and oxygen consumption rates (OCR) and extracellular acidification rates (ECAR) graphs are shown. Values were normalized to cell number generated from fluorescence intensity measurements and are represented as average of three replicates.

**Supplementary Figure 5. ONC201/TIC10 inhibits mitochondrial respiration in everolimus sensitive and resistant cells.** Cells were treated for 18 hours with indicated concentrations of ONC201/TIC10 and everolimus. Mitochondrial respiration was measured using Seahorse XF Cell Mito Stress assay and oxygen consumption rates (OCR) bar graphs are shown for basal, maximal respiration and ATP production. Values were normalized to cell number generated from fluorescence intensity measurements and are represented as average of three replicates ± SD *, P < 0.05; **, P < 0.01; ***, P < 0.001, ****, P < 0.0001.

## Star Methods

### Method details

| REAGENT or RESOURCE | SOURCE | IDENTIFIER |
| --- | --- | --- |
| Antibodies |  |  |
| BiP (C50B12) rabbit mAb | Cell Signaling | #3177 |
| pFoxO3a (Ser253) rabbit polyclonal antibody | Cell Signaling | #9466 |
| FoxO3a (D19A7) rabbit mAb | Cell Signaling | #12829 |
| pERK1/2 <sup>Thr202/Tyr204</sup> (197G2) rabbit mAb | Cell Signaling | #4377 |
| ERK1/2 (137F5) rabbit mAb | Cell Signaling | #4695 |
| pAKT <sup>Ser473</sup> (D9E) rabbit mAb | Cell Signaling | #4060 |
| AKT (C67E7) rabbit mAb | Cell Signaling | #4691 |
| pS6<sup>Ser240/244</sup> Ribosomal Protein (D68F8) rabbit mAb | Cell Signaling | #5364 |
| S6 Ribosomal Protein (54D2) mouse mAb | Cell Signaling | #2317 |
| β-actin (13E5) rabbit mAb | Cell Signaling | #4970 |
| β-actin (8H10D10) mouse mAb | Cell Signaling | #3700 |
| CREB-2/ATF-4 (B-3) mouse mAb | Santa Cruz Biotechnology | sc-390063 |
| mtTFA (TFAM) (C-9) mouse mAb | Santa Cruz Biotechnology | sc-376672 |
| TRAIL (D-3) mouse mAb | Santa Cruz Biotechnology | sc-8440 |
| CHOP (9C8) mouse mAb | Thermo Scientific | # MA1-250 |
| TUFM rabbit polyclonal antibody | Thermo Scientific | # PA5-27511 |
| Total OXPHOS Human WB Antibody Cocktail | Abcam | ab110411 |
| Bacterial and virus strains |  |  |
| LeGO-V2 (Venus) | Weber et al., 2008 | Addgene #27340 |
| LeGO-C2 (mCherry) | Weber et al., 2008 | Addgene #27339 |
| Biological samples |  |  |
| Pleural effusion or ascites samples from patients | City of Hope | IRB # 17334 |
| Chemicals, peptides, and recombinant proteins |  |  |
| Everolimus (RAD001) | Selleckchem | # S1120 |
| ONC201/TIC10 | Selleckchem | # S7963 |
| Critical commercial assays |  |  |
| CellTiter-Glo Luminescent Cell Viability Assay | Promega | G7573 |
| CellTiter-Glo 3D Cell Viability Assay | Promega | G9682 |
| EasySep CD45 Depletion Kit II | StemCell Technologies | # 17898 |
| EasySep Dead Cell Removal (Annexin V) Kit | StemCell Technologies | # 17899 |
| Agilent Seahorse XF Cell Mito Stress Test Kit | Agilent Technologies | #103015-100 |
| Experimental models: Cell lines |  |  |
| CAMA-1 | ATCC | HTB-21 |
| MCF7 | ATCC | HTB-22 |
| T47D | ATCC | HTB-133 |
| Software and algorithms |  |  |
| GraphPad Prism 9.3.1 | GraphPad |  |
| Gen5 3.05 | Biotek Instruments |  |
| Seahorse Wave Desktop Software 2.6 | Agilent Technologies |  |

### Cell culture and reagents

CAMA-1 and MCF7 breast cancer cell lines were maintained in DMEM supplemented with 10% FBS and antibiotic–antimycotic solution. T47D breast cancer cell line was maintained in RPMI supplemented with 10% FBS+ and antibiotic–antimycotic solution. Cells were regularly tested for mycoplasma contamination using commercially available Mycoplasma detection kit (Myco Alert kit, Lonza). Cell lines were authenticated using STR profiling (Laragen, Inc). Everolimus (RAD001) and ONC201/TIC10 were obtained from Selleckchem and dissolved in DMSO.

Everolimus resistant cell lines were generated by long-term culture of parental cell lines in the continuous presence of 100nM everolimus (MCF7 and T47D) or 50nM everolimus (CAMA-1) with fresh media and drug replenished every 3 days, until resistance developed (6-12 months). Resistance to everolimus was confirmed by the difference in the drug dose-response in comparison with the parental cells, measured using CellTiter-Glo Luminescent Cell Viability Assay (Promega Corporation). Everolimus-resistant cells were further maintained in complete culture medium supplemented with 50nM everolimus (CAMA-1) or 100nM everolimus (MCF7 and T47D). Sensitive and resistant cells were labeled with Venus (LeGO-V2) and mCherry (LeGO-C2) fluorescent proteins respectively as described previously (Grolmusz et al., 2020; Weber et al., 2008).

### Malignant fluid collection and primary cancer cell isolation

Malignant fluids were collected from five breast cancer patients by paracentesis (Patient # 1, 2, 4, and 5) or thoracentesis (Patient # 3) under informed consent and ethical compliance under Institutional Review Board (IRB) # 07047 and 17334 at City of Hope. Upon collection of malignant fluid, cells were pelleted at 500 x g for 5 minutes, at room temperature. Red blood cell (RBC) contamination was then removed by lysis in Tris-ammonium chloride buffer (17 mM Tris, pH 7.4, 135 mM ammonium chloride) as previously described (Nath et al., 2021) or by magnetic depletion using EasySep RBC Depletion Reagent (StemCell Technologies) diluting cell pellet in 2% FBS in PBS and incubated with 50 μL Depletion Reagent/mL for 5 minutes at room temperature, followed by two rounds of incubation on EasyEight magnet (StemCell Technologies) for 5 minutes. Primary breast cancer cells were then purified by magnetic depletion using the EasySep Dead Cell Removal (Annexin V) Kit and EasySep CD45 Depletion Kit II (StemCell Technologies) to remove dead cells and immune cells respectively as previously described (Nath et al., 2021).

### Cell proliferation

For 2D experiments, cells were plated in 384-well plates and treated with increasing concentration of everolimus (0-1μM) or ONC201/TIC10 (0-10μM) or combination ONC201/TIC10 (0-10μM) with everolimus (1nM or 100nM) for 4 days and viability was measured using CellTiter-Glo Luminescent Cell Viability Assay (Promega Corporation). Results were normalized to DMSO control treatment for each cell line. IC50 was determined using GraphPad Prism 9.3.1 software.

For 3D experiments, cells were plated in 96-well round-bottom ultra-low attachment spheroid microplate (Corning) at a density of 2.000 cells per well (for CAMA-1 cells) or 5.000 cells per well (for MCF7 and T47D cells). Spheroids were treated with drugs as indicated for up to 18 days with imaging and media change every 3-4 days. Imaging was performed using Cytation 5 imager (Biotek Instruments) gathering signal intensity from brightfield, YFP (for Venus fluorescence) and Texas Red (for mCherry fluorescence) channels. Raw data processing and image analysis were performed using Gen5 3.05 software as previously described (Grolmusz et al., 2020). Briefly, stitching of 2X2 montage images and Z-projection using focus stacking was performed on raw images followed by spheroid area analysis. Whole spheroid area and fluorescence intensity measurements of each cell line are integrated into a fitted growth equation, and cell counts for each cell line were produced from fluorescence intensities relative to spheroid size. Experiments were performed trice in triplicates and representative image is shown. Bliss Interaction Index was calculated as previously described (Soldi et al., 2013) with I >1 showing synergy and I=1 showing additivity. For 2D assay Bliss Interaction Index was calculated from luminescence values and for 3D studies from fluorescence values.

For 3D experiments with primary cells, cells were plated in 96-well round-bottom ultra-low attachment spheroid microplate (Corning) at a density of 20.000 cells per well in Renaissance Essential Tumor Medium (Cellaria) supplemented with RETM Supplement (Cellaria), 10% FBS, 25 ng/mL cholera toxin (Sigma-Aldrich) and antibiotic–antimycotic solution. After 2 days that organoid structures were formed, spheroids were treated with drugs as indicated for 4 days. Brightfield imaging was performed using Cytation 5 imager (Biotek Instruments) before treatment and after 4 days of treatment. Viability was measured using CellTiter-Glo 3D Cell Viability Assay (Promega Corporation). Results were normalized to DMSO control. Experiments were performed in triplicates and representative image is shown.

### Western blot

For immunoblot analysis cells were washed with PBS and lysed on ice in ice-cold RIPA buffer (Thermo Scientific) supplemented with protease and phosphatase inhibitor cocktail (Thermo Scientific). The protein concentration in the lysates was determined using BCA (Pierce). Equal amounts of total protein were separated by SDS-PAGE and 4% to 20% Tris-Glycine Gel (Bio-Rad), and they were transferred to nitrocellulose membranes (Thermo Scientific) according to standard protocols. Membranes were immunoblotted overnight with antibodies against BiP (C50B12), pFoxO3a (Ser253), FoxO3a (D19A7), pERK1/2 ^Thr202/Tyr204^ (197G2), ERK1/2 (137F5), pAKT ^Ser473^ (D9E), AKT (C67E7), pS6^Ser240/244^ Ribosomal Protein (D68F8), S6 Ribosomal Protein (54D2), β-actin (13E5), β-actin (8H10D10) from Cell Signaling, CREB-2/ATF-4 (B-3), mtTFA (TFAM) (C-9), TRAIL (D-3) from Santa Cruz Biotechnology, CHOP (9C8) and TUFM from Thermo Scientific. For the OXPHOS complexes detection, equal amounts of total protein were separated by SDS–PAGE and 16.5% Tris-Tricine Gel (Bio-Rad) and were transferred using the iBlot^™^ Dry Blotting system (Invitrogen) according to manufacturers’ instructions. Membranes were immunoblotted overnight with total OXPHOS Human WB Antibody Cocktail (ab110411) from Abcam that contains 5 mAbs, against Complex I subunit NDUFB8, Complex II subunit 30kDa, Complex III subunit Core 2, Complex IV subunit II, and ATP synthase subunit alpha. Experiments were performed thrice, and representative image is shown.

### Extracellular Flux Analysis

Mitochondrial respiration was determined using Agilent Seahorse XF Cell Mito Stress Test Kit according to the manufacturer’s instructions. Cells were plated in XF96 Cell Culture Microplate (Agilent Technologies Inc.) coated with Cell-Tak adhesive (Corning) at a density of 15.000 cells per well. 6 hours later cells were treated with drugs as indicated for 18 hours. Before running the assay, cell culture media were replaced with Seahorse XF assay media supplemented with 10mM XF Glucose, 1mM XF Pyruvate and 2mM XF L-Glutamine and the plate was incubated in a 37^0^C incubator without CO_2_ for 1 hour. The oxygen consumption rate (OCR) was measured by XFe96 extracellular flux analyzer (Agilent Technologies) with sequential injection of 1.5 μM oligomycin A, 1 μM FCCP, and 0.5 μM rotenone/antimycin A. For normalization, imaging was performed using Cytation 5 imager (Biotek Instruments) gathering signal intensity from brightfield, YFP (for Venus fluorescence) and Texas Red (for mCherry fluorescence) channels. Raw data processing and image analysis were performed using Gen5 3.05 software. Briefly, Gen5 3.05 software detected and generated cell counts and fluorescence intensity measurements respective for each cell line within each well. Data analysis was performed using Seahorse Wave Desktop Software 2.6. OCR was normalized to cell number. Experiments were performed trice in triplicates and representative image is shown.

### Gene expression data and pre-processing

Microarray expression data from a neoadjuvant clinical trial of ER+ breast cancer patients on everolimus were retrieved from NCBI GEO (accession number GSE119262). https://www.ncbi.nlm.nih.gov/geo/query/acc.cgi?acc=GSE119262. The tumor samples were profiled using Illumina HumanRef-8 v2 Expression BeadChips and quantile normalized using BeadArray as described in Sabine et al., 2010. The normalized gene expression matrix was aggregated by averaging expression levels of probes mapping to the same gene and by averaging the expression levels of technical replicates of the samples. RNA-seq data from MDA-MB-231 breast cancer cell lines treated with ONC201 were retrieved from NCBI GEO (accession number GSE212369). https://www.ncbi.nlm.nih.gov/geo/query/acc.cgi?acc=GSE212369. The samples were sequenced on the Illumina HiSeq 2500 platform, aligned to the h19 human genome reference using STAR, followed by gene level expression quantification using RSEM, as described in Greer et al., 2018.

### Pathway activity scores and analysis

Gene signatures for 4705 curated (C2) gene signatures were retrieved from the molecular signatures database (mSigDB version 7.0) (Liberzon et al., 2011). Single sample gene set enrichment scores were calculated from the normalized microarray (GSE119262) or RNA-seq (GSE212369) gene expression matrices using the R package GSVA (kcdf = ‘Gaussian’, method = ‘ssgsea’). To identify pathway signatures that change in response to everolimus treatment, we analyzed the ssGSEA scores of the 21 pairs of samples from patients with gene expression data collected before and after treatment. We split the samples into responsive and non-responsive groups and identified the pathways with significant difference in mean ssGSEA scores between pre- and post-treatment groups using a paired t-test. We investigated the changes in pathway signatures in response to ONC201/TIC10 treatment using gene expression data from MDA-MB-231 cells treated with ONC201 data collected at 0, 3-, 6-, 12- and 24-hour time points. We obtained ssGSEA scores for the pathway signatures at each time point and fit a generalized linear model (gaussian family) for each pathway signature against time. After obtaining nominal P-values of the fit for all signatures, we calculated false discovery rates to identify signatures that changed significantly over time.

### Statistical analysis

Dose-response curves were generated using GraphPad Prism 9.3.1 software and statistical comparisons were performed using two-way ANOVA. All data are presented as average values of samples, error bars correspond to standard deviation (SD). For all other experiments, graphs were generated using GraphPad Prism 9.3.1 and statistical comparisons of the results were performed using Student’s two tailed t test (*, P < 0.05; **, P < 0.01; ***, P < 0.001, ****, P < 0.0001).

