## Supplemental Figures 1-5 for "ONC201/TIC10 enhances durability of mTOR inhibitor everolimus in metastatic ER+ breast cancer"

### Supplementary Figure 1

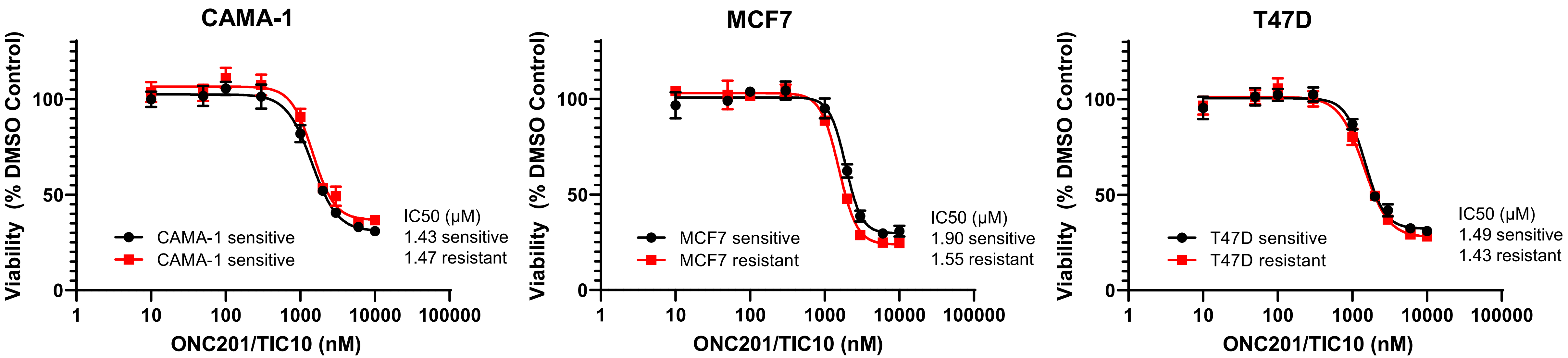

Supplementary Figure 2

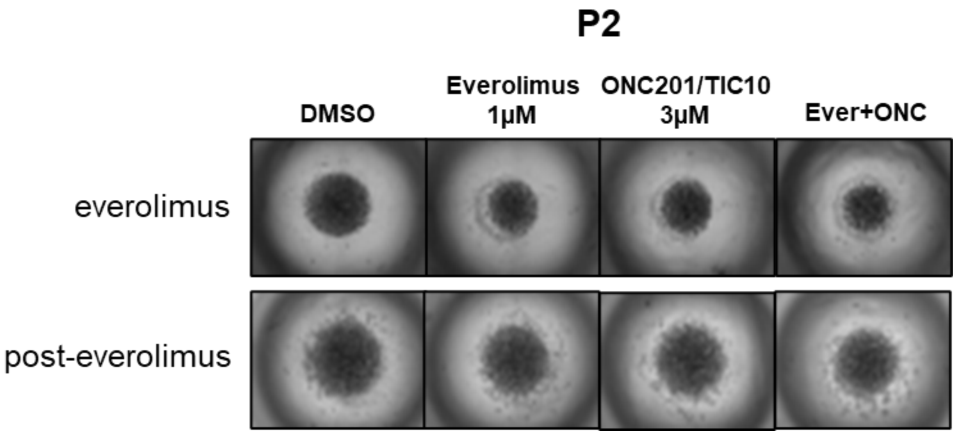

### Supplementary Figure 3

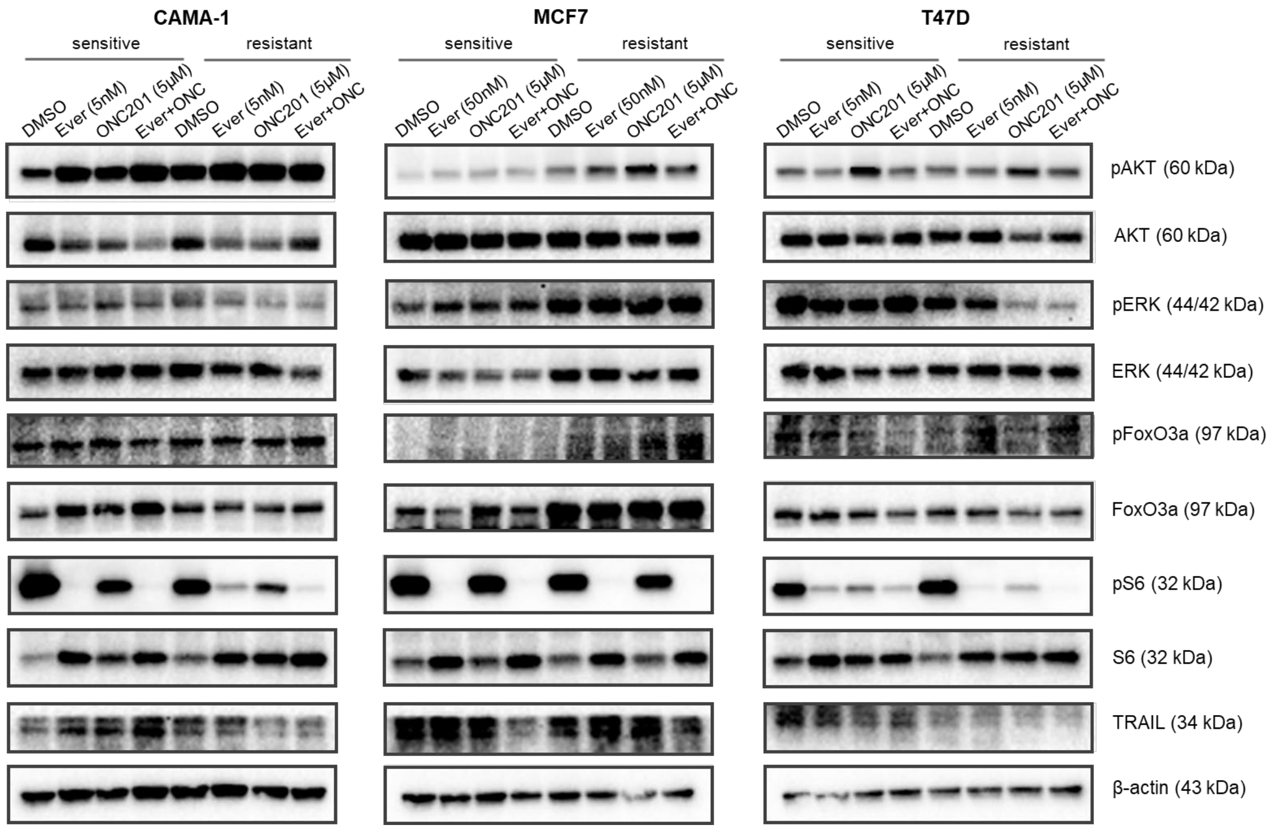

### Supplementary Figure 4

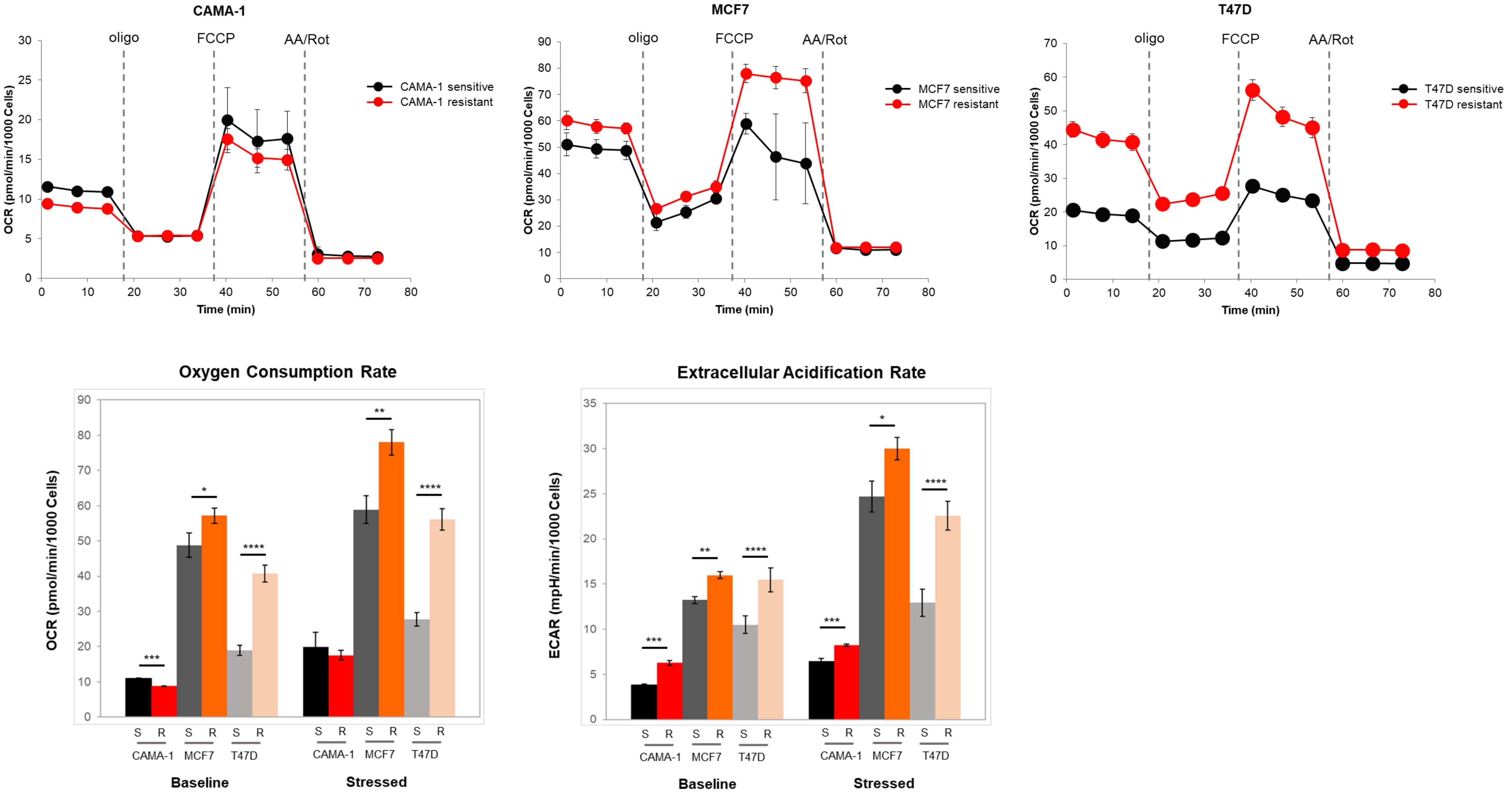

### Supplementary Figure 5

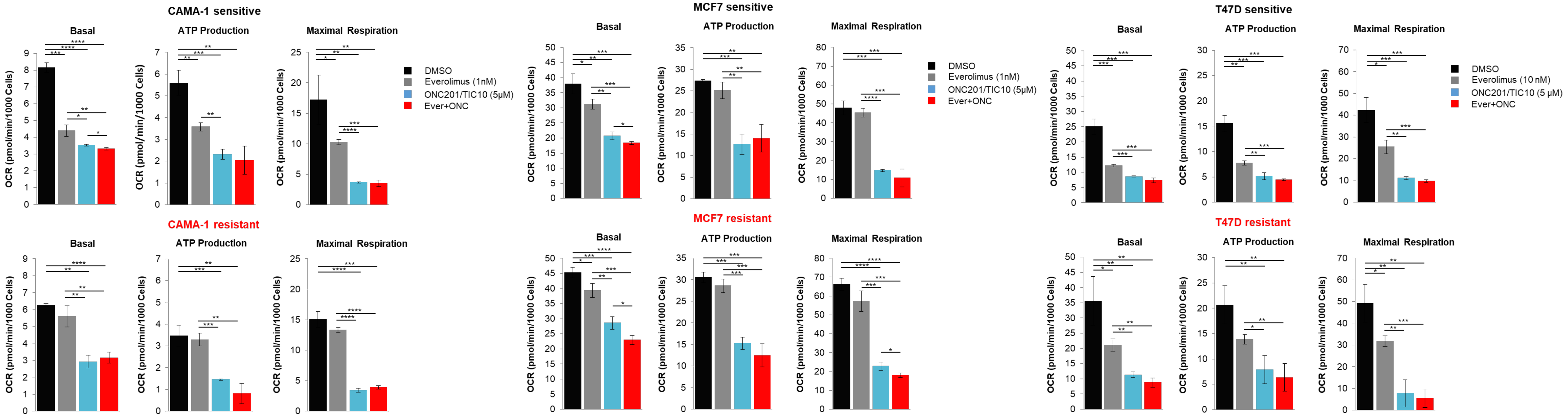
